# Negative calorie foods: An empirical examination of what is fact or fiction

**DOI:** 10.1101/586958

**Authors:** Katherine M. Buddemeyer, Ashley E. Alexander, Stephen M. Secor

## Abstract

A proposed weight loss strategy is to include in one’s diet foods that are deemed “negative calorie”. In theory, negative-calorie foods are foods for which more energy is expended in their digestion and assimilation than is consumed, thereby resulting in an energy deficit. Commonly listed negative calorie foods are characterized by a high water and fiber content and little fat. Although the existence of negative calorie foods has been largely argued against, no empirical study has fully addressed the validity of foods being negative calorie. We conducted such a study using the omnivorous lizard the bearded dragon *(Pogona vitticeps)* and celery as the tested food. Celery tops many lists of negative calorie foods due to its high fiber and low caloric content. Following their consumption of celery meals equaling in mass to 5% of their body mass, we measured from each lizard their postprandial metabolic rates to calculate specific dynamic action (SDA). Feces and urate were collected after meals to determine the energy lost to excretion. The specific energy of the celery meals, feces, and urate was determined by bomb calorimetry. Lizards lost on average 29% and 14% of meal energy to feces and urate, respectively, and an additional 33% to SDA, leaving a net gain of 24% of the meal’s energy. When considering that only a portion of fecal energy stems from the celery meal, the net gain is expectedly higher. Although this study debunks the validity of celery and other proposed foods as negative calorie, these foods will contribute to generating a negative energy budget and thus the loss of body weight.

**Summary:** This empirical study refutes the existence of negative-calorie foods; however such foods will contribute to a negative energy balance, and thus the loss of body mass.

## Introduction

From all forms of media outlets, the public is continuously bombarded by a multitude of dieting and weight loss schemes. One such dieting fad that has populated the internet and social media is a diet that consists of food considered to result in “negative calories”. In theory, these are foods for which more energy is expended in their digestion, assimilation, and nutrient storage than is gained [1–4]. Therefore their consumption results in a caloric deficit due to both the lack of net energy gained and that stored energy (i.e., fat) must therefore be utilized to fuel the completion of digestion and processing. Negative-calorie foods are generally characterized by a high fiber and water content and low caloric density [3,5,6]. Topping the well-touted lists of negative-calorie foods are celery, lettuce, grapefruit, cucumber, and broccoli [5–8].

While the conception of negative-calorie foods may be decades old, its inclusion in diets as a weight lost strategy has gained considerable interests over the past decade (largely via on-line blogs) that have promoted such foods for dieting, improved nutrition, and better heath [6,7,9]. Such diet plans have been further popularized by recent dieting books including *Foods That Cause You To Lose Weight* [10] and *The Negative Calorie Diet* [11]. The proponents of negative-calorie foods cite that in addition to generating caloric deficits, such foods possess the added benefits of boosting metabolism, controlling appetite, improving glycemic control, and cleansing your colon and liver [7,10]. Hence, the consumption of negative calorie foods results in a “win-win situation” given the multiple benefits to your health and improved weight control [6].

However, as soon as it was promoted, the validity that foods exist for which more energy is expended in their consumption than is gained was questioned. Nutritionists and physicians raised doubts of the existence of such foods citing that the cost of meal digestion and assimilation is equivalent to only 5-15% of the energy of the meal [3,12,13]. This cost refers to the accumulated energy expended on gastric acid production, gut peristalsis, enzyme synthesis and secretion, and nutrient absorption and assimilation. For humans this cost is generally referred to as diet-induced thermogenesis (DIT) whereas for other vertebrates and invertebrate it is termed specific dynamic action (SDA), the label and acronym that we will refer to in this report [14]. Therefore when accounting for SDA, it thus assumed that nearly 80-95% of the meal’s energy is still available regardless of meal type. The argument is therefore raised that even though such foods are low in caloric content, there is still a net gain in energy.

The few studies that have attempted to test the proposal that certain foods are negative-calorie have focused solely on SDA, and have produced mixed results. A study on a single participant consuming raw and liquefied celery over a 12-hr period found that the subject’s metabolic expenditure while eating either of the celery meals exceeded the energy content of the meals [15]. In contrast, Clegg and Cooper [2] calculated that the SDA of 15 female subjects following the consumption of 100 g of raw celery was less than that of the celery’s energy. The differences in these findings undoubtedly stem from the inclusion of resting metabolic rate (RMR) in the calculated energy expended in the former study and its exclusion to calculate SDA in the latter study.

In light of these studies, it has been frequently acknowledged that there is an absence of any empirical studies that have accurately tested the theory and existence of negative-calorie foods [8,12,13,16]. It should also be noted that these previous studies failed to account for the additional loss of energy in feces and urine. Given its relatively high fiber content, celery may inherently be characterized by relatively low digestive and assimilation efficiencies [17,18]. Hence, when combining the energy that is lost to DIT and to feces and urine, is it becomes theoretically more plausible for the consumption of celery to tip the balance in favor of an energy deficit [19].

We set out to address this point by employing an empirical approach that will either lend support or refute the claim that celery is a negative-calorie food. We did so by using bearded dragon *(Pogona vitticeps)*, an omnivorous lizard native to Australia. Although far removed from humans evolutionarily, bearded dragons share with humans an omnivorous diet and identical sets of mechanisms used to digest, absorb, and assimilate food [20–22]. We quantified for these lizards the energy of their celery meals, the energy expended digesting those meals, and the energy lost in feces and urate. By evaluating these energy tradeoffs, we determined bearded dragons to experience a net gain in energy from their celery meals. However, this gain is rapidly abolished by the lizard’s resting metabolism.

## Materials and methods

### Bearded dragons and their maintenance

The bearded dragon, *Pogona vitticeps* (Ahl), is a medium-sized lizard that inhabits arid desert to semi-arid woodland regions of central Australia [21,23]. It possesses a broad triangular head, rows of spiny scales along its body, and when threatened or during social display will flatten its body with males expanding their darkened throat pouch (thus their name) during social interactions [24]. Bearded dragons are naturally omnivorous and feed opportunistically on leaves, flowers, fruits, invertebrates, and small lizards [21–23]. Due to their docile nature, varied diet, and ease of captive maintenance and reproduction, the bearded dragon has become an extremely popular reptile pet [25,26]. The bearded dragons used in this study were hatched and raised in a laboratory-based colony at the University of Alabama. Lizards were housed individually or in pairs in 76-L aquariums with sand substrate, several rocks for basking, and a water dish with water available ad libitum. Light was provided by fluorescent and UVA/UVB bulbs set on a 12L:12D cycle. Room temperature was maintained at 26–29°C and humidity at 50–60%. Lizards were raised on a variety of greens (e.g., kale, collard, mustard), vegetables (e.g., carrots and squash), and calcium/vitamin dusted crickets, mealworms, and cockroaches. The nine lizards used in this study were 4-6 years of age and weighed 190.1 – 234.1 g (mean ± SE = 217.9 ± 4.9 g) at the beginning of the study.

Animal care and experimentation were conducted under an approved protocol (#14-06-0077) from the University of Alabama Institutional Animal Care and Use Committee. All efforts were made to minimize any discomfort to the bearded dragons during experimental procedures.

### Experimental procedure

We selected celery for this study because it tops many lists of reported negative-calorie foods and bearded dragons will voluntarily eat celery. We standardize meal size to 5% of lizard body mass because it is a meal size easily consumed by bearded dragons and it will generate a significant postprandial metabolic response [14,27]. Prior to metabolic and feeding trials, lizards were fasted for a minimum of 10 days to ensure that they were postabsorptive. For our colony of bearded dragons, we have observed lizards to start passing feces and urate (whitish pellet composed of uric acid) within 2-4 days after feeding. Celery was purchased at a local supermarket (Publix) and used within 24 hours of purchase. To test the claim that celery is a negative-calorie food, we quantified the gross energy of celery meals and compared that to the energy expended on celery digestion and assimilation (SDA) and the energy lost to feces and urate.

### Determination of SMR, postprandial metabolic response, and SDA

We used closed-system respirometry to quantify for each lizard their standard metabolic rate (SMR) and postprandial metabolic response [28,29]. Fasted lizards were weighed and placed into individual respirometry chambers (2.5-3 L) that were fitted with incurrent and excurrent air ports, each connected to a three-way stopcock. Respirometry chambers were placed into an environmental chamber (model DS54SD; Powers Scientific, Pipersville, Pennsylvania, USA) maintained at 30°C, with ambient air constantly pumped through the respirometry chambers. For each metabolic measurement, an initial 45-mL air sample was pulled from the excurrent port and both incurrent and excurrent ports were closed. An hour later, the excurrent port was opened and a second 45-mL sample was drawn. Air samples were pumped (75 mL min^−1^) through a column of Drierite and CO_2_ absorbent (Ascarite) into an O_2_ analyzer (S-3A/II, AEI Technologies). We calculated whole animal (mL·h^−1^) rates of oxygen consumption 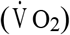 corrected for standard pressure and temperature using a modification of eq. 9 of Vleck [30].

We determined each lizard’s SMR from 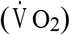 measurements while fasted. For SMR trials, 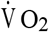 was measured in the morning (~0700) and evening (~1900) for four consecutive days. We assigned for each lizard its SMR as the mean of its two lowest measured 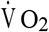 [31], Following SMR trials, lizards were returned to their cages and fed their pre-weighed celery meals. Once they had completed their meals, lizards were placed back into their respirometry chambers and measurements of 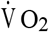 were resumed and continued at 6-h intervals for 2.5 days and thereafter at 12-h intervals for the following 2.5 days. From the postprandial measurements we determined the time span that 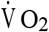 was significantly greater than SMR [14], We calculated for each lizard their SDA (kJ) by summing the extra O_2_ consumed (mL) above SMR during this time span and multiplying that total by 20.9 J. We assumed for this study that 20.9 J is expended per mL of O_2_ consumed given that the nutrient dry matter of the celery is approximately 15% protein, 5% fat, and 80% carbohydrate, and generates a respiratory quotient of 0.95 [18,32].

### Energy content of food, feces, and urate

Energy content of the celery used for each feeding trial and of the feces and urate generated from each efficiency trial was determined by bomb calorimetry. For each metabolic and efficiency trial, five sets of diced celery were dried to a constant mass at 55°C. Once dried, each sample set was reweighed (dry mass) and ground to a homogenous fine powder. A subsample of the powder from each set was placed into pre-weighed gelatin capsule (size 00, Parr Instruments, Moline, Illinois, USA), reweighed, and the capsule and powdered celery were ignited in a bomb calorimeter (model 1266; Parr Instruments, Moline, Illinois, USA) to determine total energy content. We subtracted capsule energy (19.48 kJ g^−1^ * capsule mass) from total energy to determine celery energy (kJ g^−1^ dry mass). Specific wet mass energy content of the celery (kJ g^−1^) was determined as the product of dry mass energy content and the celery’s dry mass percentage. The energy content of each ingested meal was calculated as the product of meal mass and wet mass energy content [29]. Over the course of this study, five different batches of freshly-purchased celery were used for SDA and digestive efficiency trials. For each trial, five subsets of diced celery were dried and bombed. Among samples, relative wet and dry masses were quite consistent, averaging 94.7±0.4% and 5.3±0.4%, respectively. Wet mass energy content among the samples, ranged from 0.615 to 0.933 kJ g^−1^, averaging 0. 722 ± 0.057 kJ g^−1^.

Feces and urate were collected from lizards housed individually in 76-L glass aquariums lined with laboratory countertop paper (VWR, Radnor, PA, USA) with the non-absorbent side facing upward. Prior to feeding, we emptied their large intestine by gavaging with water which removed residual feces and urate. Once fed, lizards were placed in their respective aquarium and checked twice a day for any deposited feces or urate. Any feces and/or urate found were removed, placed in individual drying trays, and dried to a constant mass at 55°C. After one week, their large intestine was gavaged of any residual feces and urate, which was also dried. For each lizard, we combined separately their feces and urate collected over the one-week period. Feces and urate resulting from these trials were dried, weighed, and bombed in gelatin capsules. After subtracting capsule energy, we calculated for each lizard the total energy of their feces and urate.

### Statistical analyses

We employed a repeated-measures analysis of variance (ANOVA) to demonstrate the statistical effects of sampling time (pre-and post-feeding) on metabolic rates. In conjunction with the ANOVA, we undertook pairwise mean comparisons (Tukey) in order to identify the post-feeding time point that lizard metabolic rates returned to values that did not differ significantly from prefeeding rates, therefore determining the duration of significantly elevated postprandial metabolism. We report means as mean ± 1SE.

## Results

### The cost of meal digestion

Fasted bearded dragons housed in darkened respirometry chambers at 30°C experienced a standard metabolic rate (SMR) that averaged 6.68 ± 0.33 mL O_2_ h^−1^ (0.030 ± 0.02 mL O_2_ g^−1^ h^−1^) (Table 1). Feeding induced a significant increase (P < 0.0001) in metabolic rate that peaked for these lizards at 12 – 24 hours postfeeding, at values that averaged 62 ± 7 % greater than SMR (Fig. 1, Table 1). Lizards maintained significantly elevated rates of metabolism for up to 3 days postfeeding. Calculated over this duration, lizards expended on average 2.64 ± 0.22 kJ digesting and assimilating their celery meals. This expenditure was equivalent to 33.1 ± 2.4% of the energy of the ingested celery (i.e., SDA coefficient [14]) (Table 1).

**Figure 1.**
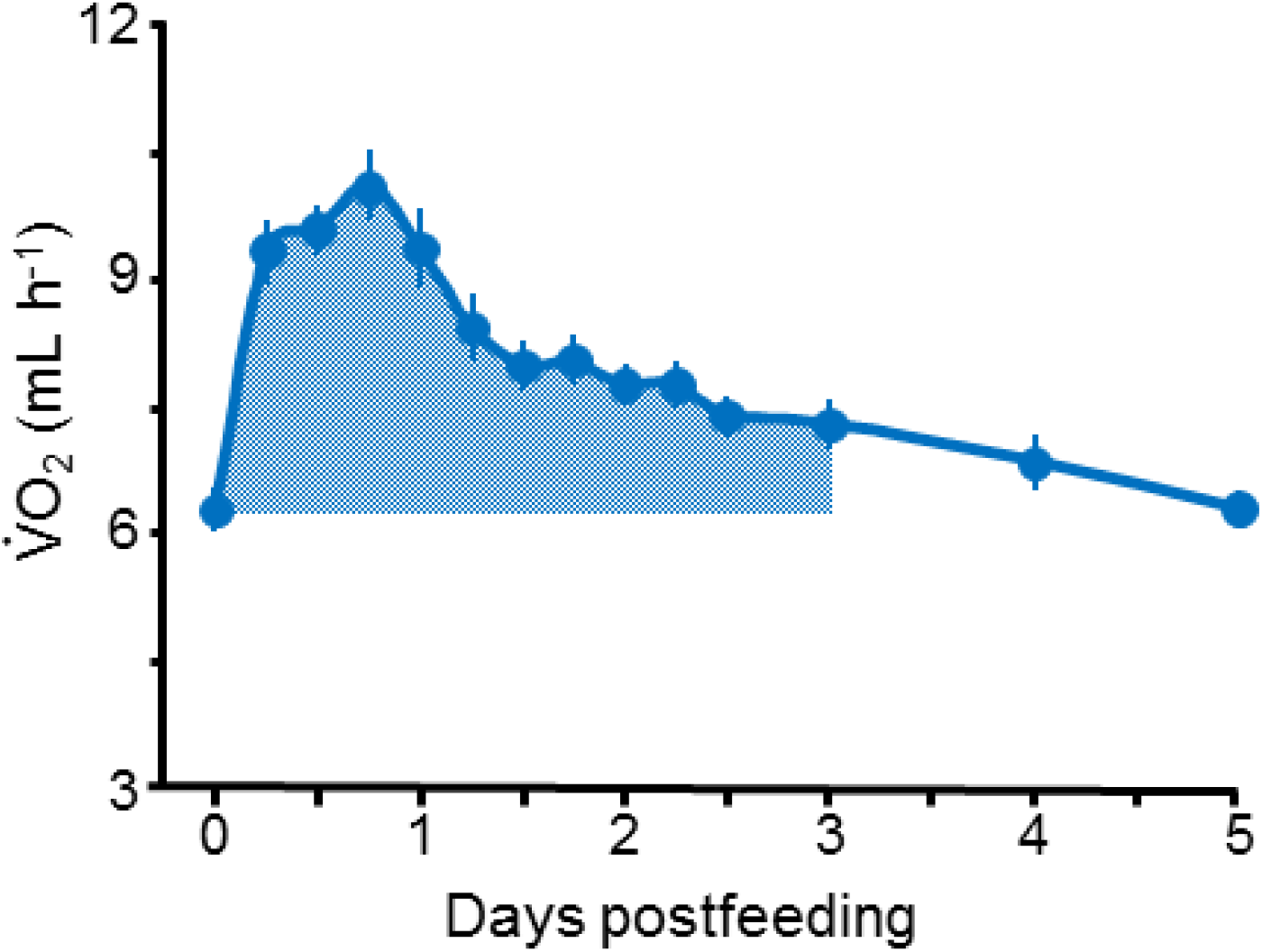
Postprandial profile of oxygen consumption and accumulative SDA. Postprandial profile of oxygen consumption (mean and SE) and accumulative SDA for nine adult bearded dragons for five days after consuming a meal of diced raw celery equivalent in mass to 5% of lizard mass. Oxygen consumption rates peaked at 24 hours postfeeding at a mean value that was 62% greater than standard metabolic rate (time = 0). The shaded area represents the extra oxygen consumed above standard metabolic rate for the three-day duration of elevated metabolism from which SDA was quantified.

**Table 1.**
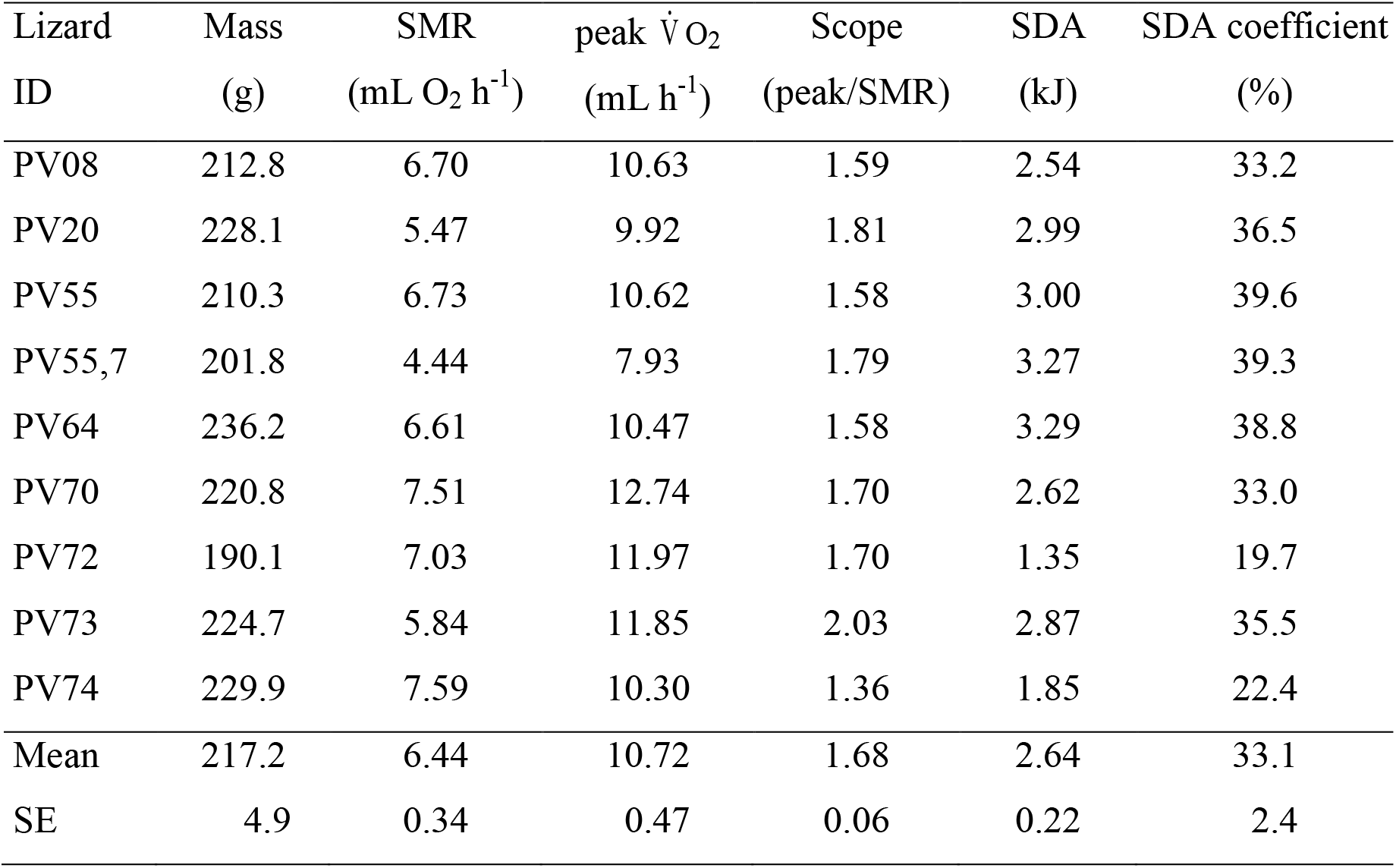
Body mass, standard metabolic rate (SMR), peak 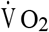, scope of peak 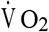, specific dynamic action(SDA), and SDA coefficient of nine adult beaded dragons *(Pogona vitticeps)* that had consumed celery meals equaling in mass to 5% of lizard body mass.

| Lizard ID | Mass (g) | SMR (mL O<sub>2</sub> h<sup>-1</sup>) | peak $\dot{V}O_2$ (mL h<sup>-1</sup>) | Scope (peak/SMR) | SDA (kJ) | SDA coefficient (%) |
| --- | --- | --- | --- | --- | --- | --- |
| PV08 | 212.8 | 6.70 | 10.63 | 1.59 | 2.54 | 33.2 |
| PV20 | 228.1 | 5.47 | 9.92 | 1.81 | 2.99 | 36.5 |
| PV55 | 210.3 | 6.73 | 10.62 | 1.58 | 3.00 | 39.6 |
| PV55,7 | 201.8 | 4.44 | 7.93 | 1.79 | 3.27 | 39.3 |
| PV64 | 236.2 | 6.61 | 10.47 | 1.58 | 3.29 | 38.8 |
| PV70 | 220.8 | 7.51 | 12.74 | 1.70 | 2.62 | 33.0 |
| PV72 | 190.1 | 7.03 | 11.97 | 1.70 | 1.35 | 19.7 |
| PV73 | 224.7 | 5.84 | 11.85 | 2.03 | 2.87 | 35.5 |
| PV74 | 229.9 | 7.59 | 10.30 | 1.36 | 1.85 | 22.4 |
| Mean | 217.2 | 6.44 | 10.72 | 1.68 | 2.64 | 33.1 |
| SE | 4.9 | 0.34 | 0.47 | 0.06 | 0.22 | 2.4 |

### Energy lost to feces and urate

Feces and urate began to appear in the cages within 2 days after feeding. Over the one-week period, lizards on average produced 0.14 ± 0.02 g dry of feces and 0.10 ± 0.01 g dry of urate (Table 2). Bomb calorimetry determined that the dry mass-specific energy content of feces and urate averaged 16.8 ± 0.6 and 11.1 ± 0.4 kJ g^−1^, respectively (Table 2). Total energy of feces and urate produced from the single celery meal averaged 2.29 ± 0.33 and 1.06 ± 0.13 kJ, respectively (Table 2).

**Table 2.** Dry mass, specific energy, and total energy of feces and urate produced by bearded dragons *(Pogona vitticeps)* within one week after consuming celery meals equaling 5% of body mass.

| Lizard ID | Body mass<br>(g) | Meal mass<br>(g) | Feces mass<br>(g) | Feces specific<br>energy (kJ g<sup>-1</sup>) | Feces energy<br>(kJ) | Urate mass<br>(g) | Urate specific<br>energy (kJ g<sup>-1</sup>) | Urate energy<br>(kJ) |
| --- | --- | --- | --- | --- | --- | --- | --- | --- |
| PV08 | 206.8 | 10.34 | 0.07 | 17.00 | 1.19 | 0.10 | 11.00 | 1.10 |
| PV20 | 229.7 | 11.48 | 0.15 | 16.41 | 2.46 | 0.08 | 11.52 | 0.92 |
| PV55 | 209.4 | 10.47 | 0.07 | 14.52 | 1.02 | 0.13 | 9.12 | 1.19 |
| PV55,7 | 202.4 | 10.12 | 0.08 | 19.64 | 1.57 | 0.07 | 10.87 | 0.76 |
| PV64 | 231.2 | 11.56 | 0.13 | 18.30 | 2.38 | 0.09 | 12.65 | 1.14 |
| PV70 | 224.0 | 11.20 | 0.19 | 14.30 | 2.66 | 0.13 | 9.79 | 1.27 |
| PV72 | 196.2 | 9.08 | 0.15 | 17.23 | 2.59 | 0.14 | 11.00 | 1.54 |
| PV73 | 223.5 | 11.20 | 0.14 | 16.45 | 2.37 | 0.12 | 11.10 | 1.37 |
| PV74 | 244.1 | 12.21 | 0.25 | 17.39 | 4.35 | 0.02 | 12.53 | 0.26 |
| Mean | 218.6 | 10.85 | 0.14 | 16.81 | 2.29 | 0.11 | 11.06 | 1.06 |
| SE | 5.2 | 0.31 | 0.02 | 0.56 | 0.33 | 0.02 | 0.38 | 0.13 |

### Net energy retained

The celery meals provided lizards with 7.83 ± 0.23 kJ of energy, of which 2.29 kJ was lost as feces, 1.06 kJ was excreted as urate, and 2.53 kJ was expended on meal digestion and assimilation, (Table 3). Therefore, the net gain in energy from these meals averaged 1.89 ± 0.17 kJ, which was equivalent to 23.4 ± 2.1% of the ingested energy. In this study, lizards retained on average nearly a quarter of their meal’s energy.

**Table 3.** Total food energy, energy lost to feces, urate and SDA, and remaining net energy (absolute and as a percentage of meal energy) for nine bearded dragons *(Pogona vitticeps)* that had consumed celery meals equaling in mass to 5% of lizard mass. Body and meal masses are the same as for Table 2.

| Lizard ID | Food energy (kJ) | Feces energy (kJ) | Urate energy (kJ) | SDA (kJ)* | Net energy gained (kJ) | Net gain as % of food energy |
| --- | --- | --- | --- | --- | --- | --- |
| PV08 | 7.47 | 1.19 | 1.10 | 2.48 | 2.70 | 36.2 |
| PV20 | 8.29 | 2.46 | 0.92 | 3.02 | 1.89 | 22.7 |
| PV55 | 7.56 | 1.02 | 1.19 | 2.99 | 2.37 | 24.9 |
| PV55,7 | 7.31 | 1.57 | 0.76 | 2.87 | 2.10 | 28.7 |
| PV64 | 8.35 | 2.38 | 1.14 | 3.24 | 1.59 | 19.0 |
| PV70 | 8.09 | 2.66 | 1.27 | 2.67 | 1.48 | 18.3 |
| PV72 | 6.56 | 2.59 | 1.54 | 1.29 | 1.14 | 17.4 |
| PV73 | 8.09 | 2.37 | 1.37 | 2.87 | 1.48 | 18.3 |
| PV74 | 8.82 | 4.35 | 0.26 | 1.97 | 2.23 | 25.3 |
| Mean | 7.83 | 2.29 | 1.06 | 2.53 | 1.89 | 23.4 |
| SE | 0.23 | 0.33 | 0.13 | 0.25 | 0.17 | 2.1 |

\*SDA was calculated as the product of meal energy and SDA coefficient of Table 1.

## Discussion

We set out to test the claim that certain foods exist for which the energy invested in their digestion and absorption, and lost through excretion exceeds the amount of assimilated energy that is gained. Thus, such foods result in a net loss of energy from the body, and therefore have been coined “negative-calorie foods”. Negative-calorie foods are characteristically high in fiber and low in energy due to their low fat and high water content [3,5,6]. We chose celery for this study because it possesses all of these characteristics and is the most cited example of a negative-calorie food [5–8]. Bearded dragons were selected for study because they are naturally omnivorous, possess a GI tract similar to that of omnivorous mammals (including humans), and can easily be studied in the lab due to their docile temperament and willingness to consume celery [20,21,26]. We partitioned the energy of the celery meals into that which is lost in SDA, feces, and urate to determine whether there was any remaining assimilated energy thus gained by the lizards. Although, the celery meals were inherently low in energy, the lizards of this study did achieve a net gain of energy from this meal.

On its face value, this empirical study debunks the claim that celery is a “negative-calorie” food, and raises doubts to the proposal that such foods do exist. However, it can be asked whether the cost of celery digestion and energy lost via excretion for the lizards approximates the equivalent cost and loss for humans.

Second, if not celery, is there a food that potentially would result in a net loss of energy if consumed? And third, which is central to its proposal for weight loss, can negative-calorie foods generate in practice a negative energy budget? In the following we will address these questions in turn.

### Costs and efficiencies of meal digestion and assimilation

The postprandial metabolic response and SDA of the *P. vitticeps* of this study are within the range observed for other lizard studies. For lizards feeding on insects, metabolic rates increase by 30 – 340%, and generate an SDA equivalent to 5 – 21% of meal energy [14]. For the only previously published vegetable diet study for lizards, the herbivorous *Angolosaurus skoogi* experienced a 78% increase in metabolic rate following the consumption of carrot meals equaling 7% of body mass [33]. Among mammalian herbivores (e.g., camel, deer, and horse) feeding on straw or hay, metabolic rates increase by 40–100% [14]. Humans tend to exhibit a relatively modest postprandial response, given that experimental mix-nutrient liquid or food diets (~1% of body mass) generate only a 20–40% increase in metabolic rate with SDA equivalent to 7–13% of meal energy [14]. To test the validity of celery as a negative-calorie food, Clegg and Cooper [2] measured the postprandial metabolic response of fifteen female subjects that had each consumed 100 g of celery (16 kcal). Subjects experienced a 33% increase in metabolic rate and generated an SDA (13.8 kcal) that equaled 86% of ingested meal energy.

Although there are noted differences in relative meal size and the SDA response between the lizards of this study and any human study, the magnitude of the metabolic increase and the profile of postprandial metabolism are similar [14]. The lizards consumed meals substantially larger, relative to body mass, than humans, and are digesting at a lower body temperature. Therefore the lizards experienced a higher postprandial peak and much longer duration of the metabolic response attributed to meal digestion and assimilation. The seemingly high cost (relative to meal energy) of digesting celery for lizards (this study) and humans [2] stems from the modest amount of energy in the celery meals. When calculated against a low meal energy, any metabolic effort will generate a higher relative cost.

It should be noted that the cost of chewing is neither calculated nor incorporated as a component of DIT or SMR [14]. Lizards and humans expend energy masticating raw celery and thus this is an additional cost that reduces the net gain of assimilated energy. For adult human subjects, the chewing of gum resulted in an increase in metabolism of 46 kJ per hour [34]. It took eight women (coauthor AA and students in the lab of SS, mean age of 21.3 years) an average of 5.4 minutes to chew (~400 chews) and swallow 100 g (four intact pieces) of raw celery. Therefore the subjects of the Clegg and Cooper [2] study potentially expended an additional 4 kJ chewing the 100 g of celery that was consumed. Although not accounted for by Clegg and Cooper [2], this added cost erases the net gain assumed in that study.

The Clegg and Cooper [2] study as well as the numerous discussions on the legitimacy of negative-calorie foods have focused only on the cost of digestion and assimilation without considering the efficiency at which these food items are digested. Humans consume foods that are easily digested and absorbed due to their highly processed nature and that they generally lack difficult to digest or non-digestable components (e.g., bone, hair, exoskeleton). What is not absorbed in the small intestine (e.g., fiber) is acted upon with in the large intestine by resident microbes (chiefly fermentation of residual carbohydrates) leaving any remaining remnants of the meal to be voided in feces. A traditional approach to quantify the efficiency of digestion has been to subtract fecal energy from meal energy and divide by meal energy [35]. The resulting “digestive efficiency” provides a metric by which comparisons can be made on the relative absorbed gain (or loss) of a meal’s energy. Taking this one step further and also subtracting the energy lost in urine (urate) before dividing by meal energy provides an efficiency index of gained assimilated energy from any meal (“assimilation efficiency”). However, there is an inherent error to this approach because it assumes that all fecal energy is derived from the undigested remnants of the meals.

Following the removal of water (~75% of fecal mass), the remaining solids of feces are in fact dominated by bacteria and other microbes (e.g., fungi, virus, protists) and sloughed intestinal epithelial cells, comprising collectively as much as 75% of fecal dry mass [17,36]. To acknowledge the inherent inaccuracy of these calculated efficiencies, they are generally reported as “apparent digestive efficiency (ADE)” and “apparent assimilation efficiency (AAE)” [37,38]. The remaining 25-40% of feces includes lipids (e.g., bacterial produced short-chain fatty acids) and undigested meal fiber. High fiber diets generate more fecal matter that is of a larger percentage of undigested fiber (relative to bacteria and sloughed cells) [17]. Celery is roughly 40% fiber (dry mass) and for the lizards, as well as for humans, a portion of that fiber is undoubtedly excreted in the feces [18].

For a back-of-the-envelope calculation, if we assume (on the high side) that lizard feces are 40% undigested fiber (roughly 30% of the ingested fiber) and add to that the total energy lost in urate (1.06 kJ) and SDA (2.60 kJ), then lizards achieve an assimilated gain of 39.9% of meal energy. This calculated gain is 70% higher than that calculated previously in the Results (Table 3), and theoretically is more accurate. How might this translate to humans digesting celery? For the only human study to assess the cost of celery digestion [2], the projected excreted energy of undigested fiber would most likely exceed the small gain that followed the subtraction of SDA, and the cost of chewing. For this study it is realistic to conclude that the participants did not gain any net calories from their celery meal. However, this study did report an extremely high SDA relative to meal energy compared to other human studies (Secor, 2009).

### Are there negative calorie foods?

For obvious reasons (low calorie, high fiber), celery has been the focus of the only empirical studies to examine the validity of negative-calorie foods [2,15, this study]. While the jury is out on whether for humans the eating of raw celery would result in no net gain of calories; are there other foods that do generate no or negative caloric gain? In absence of determining this for other foods by empirically quantifying the energy lost via feces, urate, and SDA, we can address this question by employing several assumptions of energy loss for each food. First, we set SDA equivalent to 25% of meal energy. While this coefficient is substantially lower than that calculated by Clegg and Cooper [2], it is two to three times greater than that calculated for the majority of human studies, plus it can also account for the cost of chewing [14,34]. Second, the loss of energy in urine is set at 5% of meal energy, which is similar to the loss noted elsewhere [35,37,38]. And third, that the energy lost in feces is 30% of fiber energy. For an additional nine food items commonly listed as negative calorie, the consumption of each (based on these assumptions) results in a net energy gain of roughly 64% of the ingested energy (Table 4). Even if all of fiber energy is lost in the feces (highly unlikely), energy continues to be gained from these foods (~49% on average). Double the loss to SDA and urine along with all fiber energy excreted, they continue to gain energy (~19% on average). As an exercise in budgeting energy, these calculations echo the opinions and discussions of nutritionists, trainers, physicians, and bloggers whom have debunked the existence of negative calorie foods [5,13,16,19].

**Table 4.** Tabulated for ten commonly listed negative-calorie foods is their percent water content, total energy per 100 g, total energy partitioned for carbohydrates, fiber, protein and fat, predicted SDA and energy loss in urine and feces, net gain of energy, and net gain as a percent of total ingested energy.

| Food | Celery | Broccoli | Apple | Carrot | Grapefruit | Tomato | Cucumber | Watermelon | Green<br>leaf<br>lettuce | Blueberries |
| --- | --- | --- | --- | --- | --- | --- | --- | --- | --- | --- |
| % water | 95.4 | 89.3 | 85.6 | 88.3 | 88.1 | 94.5 | 96.7 | 91.5 | 95.0 | 84.2 |
| kJ*/100 g | 71.5 | 182.4 | 254.7 | 195.0 | 207.2 | 92.3 | 55.0 | 149.9 | 81.0 | 281.7 |
| Carbs (kJ) | 52.2 | 116.7 | 242.7 | 168.4 | 187.3 | 68.4 | 38.0 | 132.7 | 50.4 | 254.6 |
| Fiber (kJ) | 28.2 | 45.8 | 42.3 | 49.3 | 28.2 | 21.1 | 12.3 | 7.1 | 22.9 | 42.3 |
| Protein (kJ) | 12.4 | 50.7 | 4.7 | 16.7 | 13.8 | 15.8 | 10.6 | 11.0 | 24.5 | 13.3 |
| Fat (kJ) | 6.8 | 14.7 | 6.7 | 9.5 | 5.6 | 7.9 | 6.4 | 6.0 | 6.0 | 13.1 |
| SDA (kJ) | 17.9 | 45.6 | 63.7 | 48.8 | 51.8 | 23.1 | 13.8 | 37.5 | 20.3 | 70.4 |
| Loss in urine<br>(kJ) | 3.6 | 9.1 | 12.7 | 9.8 | 10.4 | 4.6 | 2.8 | 7.5 | 4.1 | 14.1 |
| Loss in feces<br>(kJ)** | 8.4 | 13.7 | 12.7 | 14.8 | 8.5 | 6.3 | 3.7 | 2.1 | 6.9 | 12.7 |
| Net gain (kJ) | 41.6 | 113.9 | 165.6 | 121.7 | 136.6 | 58.3 | 34.8 | 102.8 | 49.8 | 184.5 |
| Net gain %<br>of food kJ | 58.2 | 62.5 | 65.0 | 62.4 | 65.9 | 63.1 | 63.3 | 68.6 | 61.5 | 65.5 |

\*Energy is presented in kilojoules. To convert to kilocalories, divide by 4.18. \*\*Energy lost in feces assumes that all ingested sugars, protein, and fat are absorbed and that 30% of fiber energy is lost in feces.

### Positive energy gained, negative energy budget

There are however two sides to this story. First, after accounting for the estimated energy expended on chewing, digesting, and assimilating and lost via excretion, all proposed negative-calorie foods can provide a net gain of energy. However, it is important to acknowledge that this gain only stems from the pluses and minuses specific to a meal’s digestion and assimilation and does not account for any other metabolic expenses. The other side to this story is how much this gain contributes to supporting other metabolic expenses.

The bearded dragons of this study gain theoretically as much as 3.26 kJ from their celery meal over a three-day period while at the same time expending a minimum of 9 kJ (assuming a standard metabolic rate of 0.124 kJ hr^−1^) fueling their basal metabolism and other non-digestive activities (i.e., moving in their enclosures). Whereas lizards did achieve a positive energy budget specific to digestion and assimilation, they experienced a negative energy budget over those three days (a loss of at least 6 kJ) (Fig. 2). Lizard enthusiasts would never consider celery as a stable diet for bearded dragons, and rather instead feed them higher quality vegetables and greens along with insects [26].

**Figure 2.**
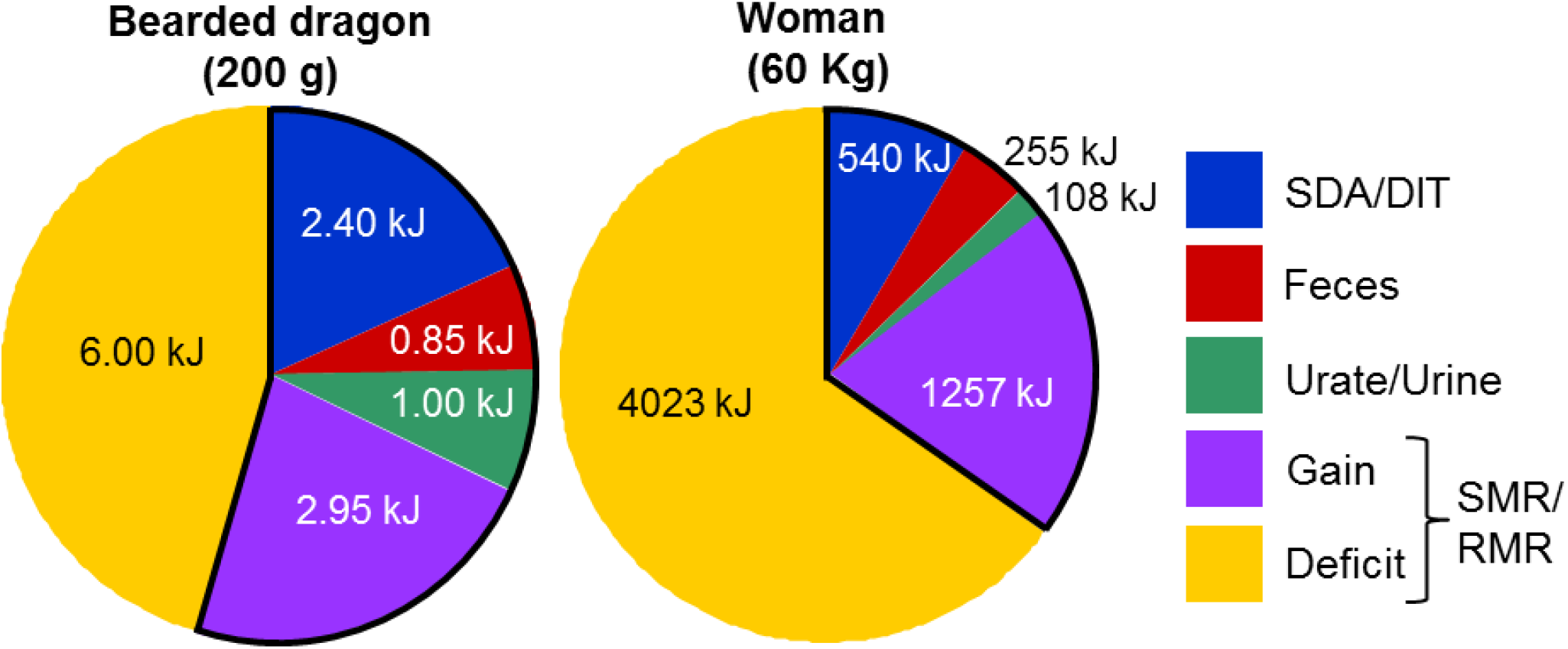
The partitioning of meal energy and energy deficit from celery meals. Pie charts illustrating the energy partitioned to SDA/DIT, feces, urate/urine, and metabolizable gain (outlined in black) for raw celery meals equaling in mass to 5% of body mass over a 3-day period for 200-gram bearded dragon and over a 1-day period for 60-kg woman. The total energy of each chart represents the predicted expenditure on standard metabolic rate (SMR, at 30C) for the bearded dragon for 3 days and on resting metabolic rate (RMR) for a woman for 1 day. The noted deficit for each is the amount of additional energy needed to fuel SMR or RMR for 3 days and 1 day, respectively, for lizards and women, beyond that gained from the celery meal (deficit + gain = SMR or RMR). The fuel to cover this deficit undoubtedly originates from endogenous body stores.

The same is undoubtedly true for humans. Those foods touted as negative calorie do generate a net energy gain; however this gain is quickly abolished by the body’s own basal rate of metabolism. Consider that a 60 kg woman possesses a resting metabolic rate of approximately 220 kJ/h [39]. If we assume that SDA is equivalent to 25% of meal energy (including the cost of chewing) and the woman loses 5% of meal energy in her urine and 30% of fiber energy in her feces, then a celery meal of 5% of body mass (3 kg) would only provide the fuel to cover a little less than six hours of her resting metabolism (Fig. 2). Cut that time in half if she is active. Following these same assumptions, this woman would need to consume daily 9100 kJ or 12.6 kg (~28 lbs) of raw celery (given the loss to SDA, feces, and urine) to fuel her resting metabolism for that day. It is unlikely that anyone would maintain a daily diet of 12.6 kg of raw celery, or 9 kg of tomatoes, or even 4.3 kg of raw carrots just to fuel their minimal metabolic needs.

The central aim of the majority of weight loss programs is to achieve a negative energy balance; in concept, one’s daily energy expenditure (DEE) exceeds their daily metabolizable energy intake (MEI; meal energy minus energy lost in feces and urine). Hence, DEE must be supplemented by the catabolism of endogenous energy stores, chiefly the body’s stores of fat. This can be accomplished by increasing expenditure and/or decreasing intake such that DEE > MEI. Increasing the relative proportion of the diet that includes low calorie, high fiber foods serve the right side of the equation, reducing MEI. This is the game plan that dominates those diet plans that campaign for the inclusion of proposed negative calorie foods as a surefire means to burn fat and lose weight [5–8,10,11].

In this study we empirically tested the theory that a low-calorie, high fiber food would generate an energy deficit due to a cost of processing that exceeds energy gain, and thus be negative caloric. Bearded dragons gained energy from their celery meals (refuting the negative calorie claim), however the energy gained contributes very modestly to fueling their DEE. Rather than labeling such foods as “negative calorie” it would be more accurate to pitch these foods as “negative budget”, the consumption of which will favor a daily negative energy budget, and hence weight loss via the catabolism of body fat.

## Acknowledgments

This study was conceived following the introduction of the concept of negative calorie foods to the senior author by his son (a sophomore and English major at the time) who proposed to conduct a human study with his friends. However, when it was explained to him the additional need to collect and bomb everyone’s feces, he declined. For assistance with this study, we thank Mimi Bach, Kellen Cowen, Tori Fields, Georgia Gamble, Ayla Jones, Mackenzie Kyler, Alexis McGraw, Zoe Nichols, Anna Reding, and Amanda Shoemaker.

## References

1. O’Connor A. The Claim: Some Foods Have Negative Calories. 2006 July 25 [cited 3 March 2019]. In: The New York Times, Health [internet]. [about 1 screen]. Available from: https://www.nytimes.com/20006/07/25/health/25real.html.

2. Clegg ME, Cooper C. Exploring the myth: Does eating celery result in negative energy balance? Proc Nutr Soc. 2012; 71: E217.

3. Wilson C. Negative-Calorie Foods” Still Count. 2016, Aug 26 [cited 3 March 2019]. In: eatright [Internet]. Chicago: Academy of Nutrition and Dietetics. [about 2 screens]. Available from: https://www.eatright.org/health/weight-loss/fad-diets/negative-calorie-foods-still-count.

4. Cespedes A. What is the Negative Calorie Diet?. 2017 Oct 20. [cited 3 March 2019]. In: workingmother.com [internet]. [about 2 screens]. Available from: https://www.workingmother.com/momlife/13527533/what-is-the-negative-calorie-diet/.

5. Dunford L. Negative Calorie Foods Are A Myth – Here’s Why. 2018, Aug 22. [cited 3 March 2019]. In: Independent.co.uk [internet]. London. Independent Digital News & Media. [about 3 screens]. Available from: https://www.independent.co.uk/life-style/food-and-drink/calorie-foods-counter-myth-healthy-eating-weight-loss-negative-a8500021.html.

6. Sengupta S. Negative Calorie Foods: You Can Eat These 11 Foods & Not Gain Weight. 2018, March 26. [cited 3 March 2019]. In: Food.ndtv.com [internet]. New Delhi Television. [about 7 screens]. Available from: https://food.ndtv.com/food-drinks/11-foods-that-burn-more-calories-than-they-contain-1679965.

7. Biswas C. Negative Calorie Foods – What Are They, How They Work, And Benefits. 2018 Nov 13. [cited 3 March 2019]. In: Stylecraze, weight loss [internet]. [about 5 screens]. Available from: https://www.stylecraze.com/articles/negative-calorie-foods-list/#gref.

8. Lall A. 7 ‘Negative-Calorie’ Foods That Help You Stay Satisfied While Dieting. 2018 Aug 16). [cited 3 March 2019]. In: First for Women [internet]. Englewood Cliffs, New Jersey [about 5 screens]. Available from: https://www.firstforwomen.com/posts/negative-calorie-foods-164691.

9. Streit L. 38 Foods That Contain Almost Zero Calories. 2018 June 11. [cited 3 March 2019]. In healthline [internet]. [about 15 screens]. Available from: www.healthline.com/nutrition/zero-calorie-foods.

10. Barnard N. Foods that cause you to lose weight, the negative calorie effect. New York: William Morrow; 2016.

11. Dispirito R. The negative calorie diet. New York: Harper Collins; 2016.

12. Nelson J. Do negative-calorie foods really exist? 2015 June 9. [cited 3 March 2019]. In: MNN.com, Food & Drink, Healthy Eating [internet]. Mother Nature Network, Narrative Content Group. [about 4 screens]. Available from: https://mnn.com/food/healthy-eating/stories/do-negative-calorie-foods-really-exist.

13. Langer A. (Diet Review) Negative Calorie Foods Don’t Exist, So Forget That Nonsense And Get On With Your Life. 2017 Aug 8. [cited 3 March 2019]. In: Abby Langer Nutrition [internet]. Toronto, Canada. [about 6 screens]. Available from: https://abbylangernutrition.com/diet-review-negative-calorie-foods-dont-exist-forget-nonsense-get-life/.

14. Secor SM. 2009. Specific dynamic action, a review of the postprandial metabolic response. J Comp Physiol. 2009; 179: 1–56.

15. Hughes T. Eating celery really DOES burn more calories than it contains. 2016 June 10. [cited 3 March 2019]. In: Daily Mail, Health [internet]. [about 2 screens]. Available from https://www.dailymail.co.uk/health/article-3636165/Eating-celery-really-DOES-burn-calories-contais.html.

16. Pike A. Mythbuster: Negative Calorie Foods. 2016 Oct 11. [cited 3 March 2019]. International Food Information Council Foundation. [internet]. [about 2 screens]. Available from: https://www.foodinsight.org/myth-zero-calorie-foods.

17. Rose C, Parker A, Jefferson B, Cartmell E. The characterization of feces and urine: A review of the literature to inform advanced treatment technology. Crit Rev Environ Sci Tech. 2015; 45: 1827–1879.

18. U.S. Department of Agriculture, Agricultural Research Service. USDA National Nutrient Database. Nutrient Data Laboratory. 2018. Available from: http://ndb.nal.usda.gov.

19. Dunning B. Negative Calorie Food Myths. 2012 Aug 7. [cited 3 March 2019]. In: skeptoid.com [internet]. [about 4 screens]. Available from: https://skeptoid.com/episodes/4322

20. Hernandez RA, Secor SM, Espinoza RE. Is a dietary jack of all trades a master of none? Adaptability of gut form and function in an omnivorous lizard. Integr Comp Biol. 2005; 45: 1011.

21. Cogger HG. Reptiles and Amphibians of Australia. 7th ed. Clayton, Australia: CSIRO Publishing; 2014

22. Oonincx DG, van Leeuwan JP, Hendriks WH, van der Poel AF. The diet of free-roaming Australian central bearded dragons *(Pogona vitticeps)*. Zoo Biol. 2015; 34: 271–277.

23. Wilson S. Australian lizards: A natural history. Clayton, Australia: CSIRO Publishing; 2013

24. Brattstrom BH. Social and thermoregulatory behavior of the bearded dragon, *Amphibolurus barbatus*. Copeia.1971; 1971: 484–497.

25. Mazorlig T. Bearded dragons (Animal plant pet care library). Neptune, New Jersey: TFH Publications, Inc; 2011.

26. De Vosjoil P, Sommella TM, Mailloux R, Donoghue S, Klingenberg RJ. The bearded dragon manual: Expert advice for keeping and caring for a healthy bearded dragon. Metuchen, New Jersey: Companion House Books; 2016.

27. Alexander AE, Buddemeyer KM, Secor SM. Testing the cooking hypothesis in human evolution. Integr Comp Biol. 2015; 55: e212.

28. Secor SM, Diamond J. Determinants of post-feeding metabolic response in Burmese pythons *(Python molurus)*. Physiol Zool. 1997; 70: 202–212.

29. Crocker-Buta SP, Secor SM. Determinants and repeatability of the specific dynamic action of the corn snake, *Pantherophis guttatus*. Comp Biochem Physiol. 2014; 169A: 60–69.

30. Vleck D. Measurement of O2 consumption, CO2 production, and water vapor production in a closed system. J Appl Physiol. 1987; 62: 2103–2106.

31. Bessler SM, Stubblefield MC, Ultsch GR, Secor SM. 2010. Determinants and modeling of specific dynamic action for the garter snake, *Thamnophis sirtalis*. Can J Zool. 2010; 88: 808–820.

32. Gessaman JA, Nagy KA. Energy metabolism: errors in gas-exchange conversion factors. Physiol Zool. 1988; 61:507–513.

33. Clarke BC, Nicolson SW. Water, energy, and electrolyte balance in captive Namib sand-dune lizards *(Angolosaurus skoogi)*. Copeia. 1994; 1994: 962–974

34. Levine J, Baukol P, Pavlidis I. 1999. The energy expended in chewing gum. New Engl J Med. 1999; 341: 2100.

35. Brody S. Bioenergetics and growth. New York: Hafner; 1945

36. Bojanova DP, Bordenstein SR. Fecal transplants: What is being transferred. Plos Biology. 2016; 14: e1002503.

37. McConnachie S, Alexander GJ. The effect of temperature on digestive and assimilation efficiency, gut passage time and appetite in an ambush foraging lizard, *Cordylus melanotus melanotus*. J Comp Physiol B. 2004; 174:99–105.

38. Cox CL, Secor SM. 2007. Determinants of energy efficiencies in juvenile Burmese pythons, *Python molurus*. Comp Biochem Physiol. 2007; 148A: 861–868.

39. Siervo M, Boschi V, Falconi C. Which REE prediction equation should we use in normal-weight, overweight and obese women? Clinical Nutrition 2003; 22: 193–204.

